# Developing machine-learning-based amyloid predictors with Cross-Beta DB

**DOI:** 10.1101/2024.02.12.579644

**Authors:** Valentin Gonay, Michael P. Dunne, Javier Caceres-Delpiano, Andrey V. Kajava

## Abstract

Due to shifts in environmental conditions, mutations, or interactions with other biomolecules, some proteins that would normally be soluble can undergo aggregation, resulting in the formation of clumps of amyloid fibrils. Understanding of this phenomenon is of paramount importance due not only to its association with various diseases (including Alzheimer’s disease), but also due to increasingly abundant evidence for its functional roles. Numerous studies have demonstrated that the propensity to form amyloids is coded by the amino acid sequence and this finding has paved the way for the development of several computational predictors of amyloidogenicity. The ultimate objective of computational methods is to accurately predict the formation of disease-related and functionally relevant amyloids that occur *in vivo*. These amyloid fibrils are known to form very specific “cross-β” structures of protein regions longer than about 15 residues. Remarkably, despite the significance of the naturally occurring amyloids, there has been a lack of datasets specifically dedicated to them. Hence, we built Cross-Beta DB, a database composed of cross-β amyloids formed in natural conditions. This database is expected to be indispensable for benchmarking amyloid predictors. We used the Cross-Beta DB to train and benchmark several such algorithms, using machine learning. The best-performing of these, the random-forest-based Cross-Beta RF Predictor, demonstrated superior performance over the other existing methods, fostering high expectations for an improved prediction of naturally occurring amyloids.

## Introduction

Proteins are often required to be soluble in order to effectively carry out their cellular functions, which typically involve interactions with other proteins or biomolecules. Numerous studies have shown that proteins can lose their solubility through a process of misfolding and aggregation (Steven et al., 2016), which not only disrupts their function, but can prevent their degradation by proteases and cause harmful accumulation within cells. Changes in solubility can be triggered by factors such as mutations, environmental changes, or interactions with other proteins, referred to as prions. In some cases, the insoluble proteins self-assemble into protease-resistant aggregates, often taking the form of amyloids. In some cases amyloids are functional, for example the RIP1/RIP3 complex in the programmed necrosis pathway is (Li et al., 2012). However they have been more widely studied in the context of diseases, typically as a buildup of amyloid fibrils in the form of amyloid plaque present in various tissues. For example, amyloidosis in brain tissue can cause neurodegenerative diseases, such as is the case with the Tau protein and Aβ peptide in Alzheimer’s disease (Miller-Thomas et al., 2016). The amyloid protofibrils form a very specific structure called the cross-β structure in which the polypeptide chains are oriented perpendicular to the fibril axis (Kajava et al., 2006). In the cross-β structure of naturally occurring amyloids, the β-strands are parallel, following the N to C-terminal direction, and are in-register (Tycko, 2011, Iadanza et al., 2018) (Figure 1A). One of the most important yet often overlooked aspects concerning naturally occurring amyloids is the minimal length of the regions that form these structures. While both short and long amyloidogenic peptides can form fibrils in vitro at high concentrations, it does not necessarily mean that these peptides will exhibit equal fibrillization efficacy under in vivo conditions. Indeed, the naturally occurring amyloid-forming proteins with amyloidogenic regions longer than 15 residues are well-documented (Kajava et al., 2010 & Iadanza et al., 2018). At the same time, it has been demonstrated that fusing short amyloidogenic peptides with soluble proteins can only induce fibrillation in vitro at high concentrations of 300–600 □M (Esteras-Chopo et al., 2005 & Guo & Eisenberg, 2008). Thus, it can be inferred that the length of amyloid-forming regions is a crucial factor, and in the majority of naturally-occurring cases, including both disease-related and functional amyloids, these regions extend beyond approximately 15 residues.

**Figure 1.**
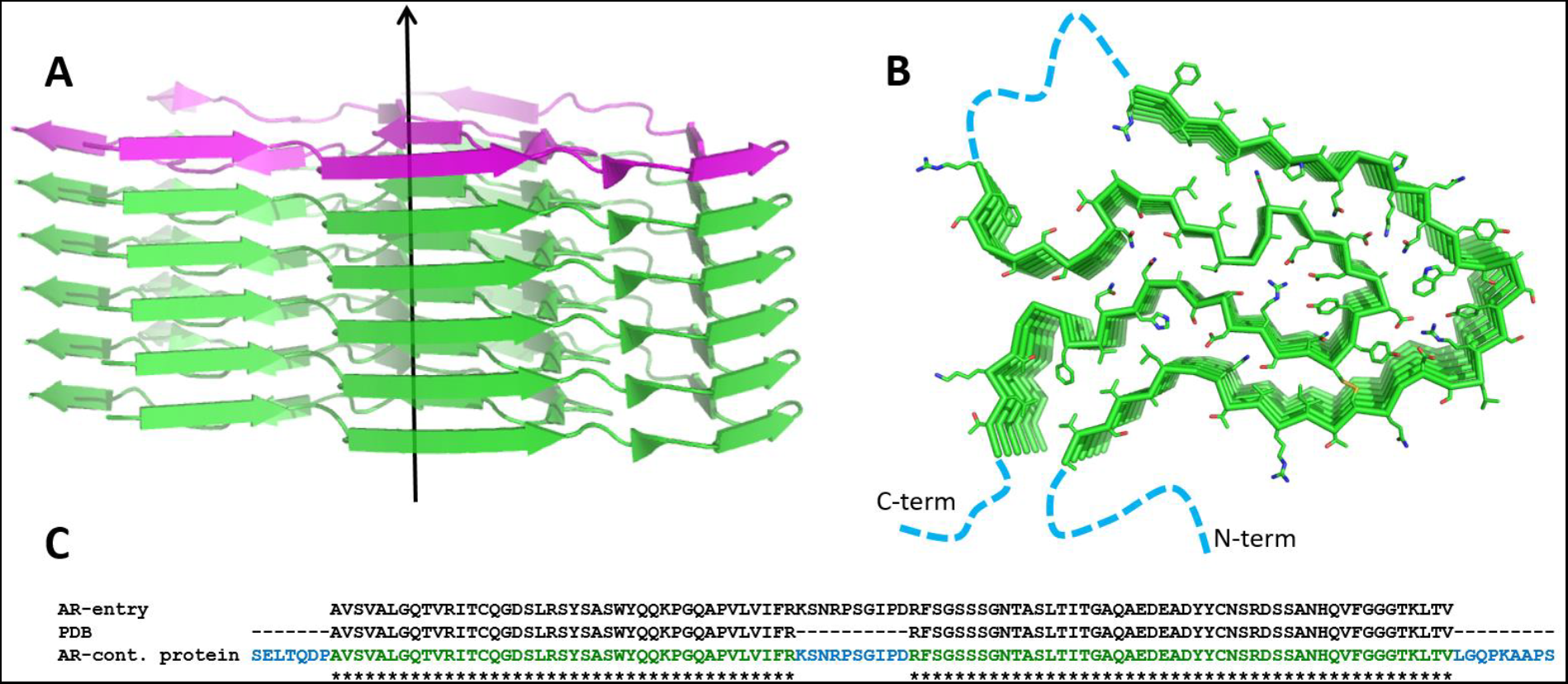
Cross-β structure of amyloid fibrils with parallel and in-register arrangement of the chains. **A**. Lateral view of a cross-b amyloid fibril formed by the immunoglobulin lambda light chain variable region [*Homo sapiens*] (PDB entry 6Z1O). The arrow shows the fibril’s axis and one of the individual chains is shown in magenta. **B**. Axial view of the amyloid fibril (6Z1O). Blue dotted lines denote regions of the protein that were not resolved in the structure. **C**. Amino acid sequence alignment of the AR-containing protein, ordered part of the fibril in accordance with its PDB entry and AR entry of Cross-Beta DB. Sequence of AR-containing protein is colored similarly to the chain shown in **B**.

Another crucial consideration is the presence of the cross-β structure in the aggregates (Figure 1A). Not all aggregates may exhibit this structure; some of them may be amorphous or alpha-helical and may not be associated with amyloids (Fink, 1998). Various techniques are available to identify cross-β structures in amyloid fibrils, with some of the most powerful methods being cryo-Electron Microscopy (cryo-EM) (Gremer et al., 2017), solid-state Nuclear Magnetic Resonance (ss-NMR) (Tycko, 2011), and fiber diffraction (usually with X‐rays, but sometimes using neutrons or electrons) (Jahn et al., 2010).

Other methods, such as Thioflavin T (ThT) and Congo-red binding assays, CD and IR spectroscopies, AFM or EM (Guo & Akhremitchev, 2006 & Chun et al., 2022), can also offer insights into the likelihood of fibrils having the cross-β structure. However, it is important to note that this information is not straightforward and requires interpretation in combination with other data. For example, ThT binds to cross-β fibrillar structures, but also to some other protein structures (Biancalana & Koide, 2010). The Congo-red (Howie, 2019 & Espargaró et al., 2020) is another dye used for the detection of amyloids. Upon binding to the cross-β amyloids, it exhibits a specific birefringence, showing green and yellow refraction under polarized light. But again, it can have the same effect when bound to the other periodic structures. As the volume of biology-related information is rapidly expanding, the significance of databases containing this data is also on the rise. The data on the amyloids is no exception to this rule.

The most complete source of information on the 3D structure of amyloids is the Protein Data Bank (PDB) (Berman et al., 2000). The other, more specialized database dedicated to the amyloid proteins is AmyPro (Varadi et al., 2018). These existing databases have their own criteria for entry selection and contain different sets of entry-related data. For example, the WALTZ-DB (Louros et al., 2020) comprises short peptides (about 6 residues), known to either form or not form amyloid fibrils under in vitro conditions.

The amyloid-related databases have become especially important for the benchmarking of amyloid predictors and training of machine learning tools. Indeed, the accuracy of prediction methods, especially those based on machine learning, depends significantly on the quality of the data used for their training. The eventual objective of such bioinformatics tools is to accurately predict naturally occurring amyloids based on the analysis of amino acid sequences. For this purpose, the presence of the database containing only the known amyloid-forming regions of the naturally-occuring cross-β amyloid fibrils is critical, however until now no such database has been developed. Here we present the Cross-Beta DB, a carefully curated database that aims to fill this gap. The construction of this database allowed us to build machine-learning-based predictors of amyloidogenicity that demonstrated superior performance over existing methods, which we also present here.

## Results and Discussion

### A. Selection of database entries

The key element of each entry in Cross-beta DBis the amyloidogenic region (AR). We selected entries using the following criteria: (i) the amyloid is naturally-occuring (functional or disease-related), (ii) AR localization within the protein is known and (iii) the amyloid structure has the cross-β arrangement.

The majority of entries of the Cross-Beta DB Release #1 were acquired from the Protein Data Bank (PDB) (January 2024 release), which contains the most detailed and up-to-date information on the atomic structures of the amyloids. To find the amyloid fibril structures, we used the following query: “*(Has Experimental Data = “Y” AND Structure Determination Methodology = “experimental” AND Polymer Entity Sequence Length >= 15) AND (Full Text = “Amyloid” AND Full Text = “fibril” AND Full Text = “Cross-Beta”)”*. From the 3072 query results, 250 candidates, showing structure of amyloid fibrils, were selected based on visual inspection of the 3D structures.

After selection, the chosen sequences were subjected to a series of processing steps. In the PDB, the N- and (or) C-terminal sequences of some entities did not belong to the amyloid structure or protruded from it in a disordered conformation. We manually removed the corresponding residues from the sequence. An example of such an entry is the fibril structure of Abeta40 peptide (PDB ID 2LMN, region G680-K687) (Paravastu et al., 2008). Some ARs also had gaps in the sequences, which correspond to unresolved fragments of the structures. An example is the Amyloid lambda 3 light chain (28-126) (PDB ID: 6Z1O) (Radamaker et al., 2021) (Figure 1B). The missing sequences of the gaps were restored from the original protein sequences found in the UniProt database (The Uniprot Consortium, 2023) by using BLAST (Altschul et al., 1990). Additionally, some entries were taken from AmyPro database, which is another large database of amyloid-forming proteins (Figure S1). First, we identified entries from the AmyPro database that were absent in the PDB by comparing the AR sequences. We then filtered out AR sequences from AmyPro that were less than 15 residues long (considered as a minimal length required to induce the formation of naturally-occuring cross-β amyloids). The AmyPro entries were then examined for the presence of the cross-β structure in the fibrils based on the literature data. Only evidence coming from experimental studies including X-ray fiber diffraction, electron diffraction via the transmission electron microscope, ss-NMR spectroscopy, Congo-red birefringence, electron microscopy, FTIR and CD spectroscopy was considered. If the X-ray fiber diffraction, electron diffraction provide direct evidence of the cross-β arrangement, the other methods allow us to draw conclusions based on the sum of the indirect data. For example, Osmotically-inducible lipoprotein B (28-44) AP00087 (DB00088) in accordance with electron microscopy (EM) forms straight and unbranched fibrils typical for the cross-β structures, CD and FTIR spectroscopy data suggest beta-conformation of the protein and Congo-Red birefringence is observed (Jarrett & Lansbury Jr, 1992). On the other hand, Lung Surfactant protein (32-57) (AmyPro ID: AP00130) was excluded because of a lack of clear evidence on the cross-β structure having only data on ThT / Congo-red binding, (Szyperski et al., 1998).

Ranges of allowed pH and temperature were the other selection criteria. We included only entries of amyloid fibrils formed between temperatures of 4°C to 40°C. Among excluded entries were the AP00030 entry of AmyPro, because its amyloid fibrils are formed at a temperature of 65°C (Fändrich et al., 2003), and AP00047, for which amyloid formation was observed only after freezing and thawing (Graether et al., 2003). As concerns pH, we considered only values of more than 5.5 and less than 9.0. The rationale behind this choice was that within the chosen range of pH amino acid side-chains should not change their electric charges. Individual glutamines become neutral below pH 4.1 and Lys above pH 10.1. At the same time, within the parallel and in register cross-beta structure, where amino acids with the same charge are in close proximity to each other, this range can be narrower. For example, it has been shown that glutamates inside of the Beta-endorphin amyloid fibril are neutral and form H-bonds with each other at pH 5.5 (Seuring et al., 2020). As concerns lysines, although the experimental data about their pKa in the cross-β structure are absent, the upper limit of pH should be less than 10.1. We have chosen the upper limit as pH 9.

The selection criteria employed for AmyPro entries have also been applied to the search for new cases of cross-β amyloids in the scientific literature. Some of 173 selected entries had the same amyloid forming sequence. They have been grouped together by using clustering based on amino acid sequence and length and are presented in our database as a single representative entry (for details see Scheme on Figure S1). As a result of the selection and exclusion criteria, the current version of the Cross-Beta DB comprises 116 entries, with 99 sourced from PDB, 10 from AmyPro, and 7 from the literature search.

### B. Cross-Beta DB entry enrichment

Starting from our set of Cross-Beta DB ARs, we expanded the entries with supplementary information consisting of 39 different variables. For example, we added the complete sequence of the AR-containing proteins. In case of frameshift or stop codon mutations located within ARs, we used the best match as the AR-containing protein sequence and preserved the author’s amino acid numbering. We also added amino acid positions of disordered (flexible) regions within the fibril 3D structure. The numbering used corresponds to the positions of the AR-containing proteins. This was the case of the amyloid fibril from a lambda light chain (PDB ID: 7NSL (Radamaker et al., 2021), 6IC3 (Radamaker et al., 2019)) or of the cardiac amyloid fibrils from immunoglobulin light chain (PDB ID: 6HUD (Swuec et al., 2019)). Some entries of our database contain information about ligands or chemical modifications associated with the PDB structures. As an example, the C-terminal residue of the islet amyloid peptide from the structure PDB (ID: 6Y1A) has an NH2-group registered in the ligand column (Roder et al., 2020). The Ser8 phosphorylated beta-amyloid 40 fibril (PDB ID: 6OC9) has 2PO phosphonate in this column (Hu et al., 2019). Beta-amyloid 42 fibrils of the PDB entry 7Q4B bind cations recorded as unknown atom or ion (UNX) (Yang et al., 2022).

For each selected entry from the PDB we downloaded the following information: Entry ID (PDB code), Entity ID (ID of polypeptide chain that form the amyloid fibril), Database Name (database with more detailed information about the protein), Accession Code(s) (in the database), Sequence, Polymer Entity Sequence Length (length of the amyloid-forming sequence), and Molecular Weight (Entity) (in kDa). In addition, because of the corrections applied to the AR sequences (for example Figure 1B, C), we re-calculated the sequence length and molecular weight of the ARs as well as their amino acid composition in the form of a vector of 21 dimensions (20 natural amino acids and ‘X’ corresponding to chemically modified amino acids). An example of a PDB entry with ‘X’ in the sequence is 7M61 (Cao et al., 2021).

ARs of some entries have mutations. For example, transthyretin (PDB ID: 6SDZ) (Schmidt et al., 2019) has a mutation V50M, transcription elongator 1 (PDB ID: 2NNT) has a mutation Y446F (Ferguson et al., 2006). The information about the mutations was recorded in a separate column.

The entries of the Cross-beta DB also have several predicted characteristics such as AR and Exposed Amyloidogenic Regions (EAR) detected by ArchCandy2.0 (Falgarone et al., 2023), disorder predictions using IUPred3 (Erdős et al., 2021) and structure prediction by ESM fold (Rives et al., 2021).

Finally, we grouped the entries with identical AR sequences in one representative entry recording the other PDB IDs in the ‘Other ID’ column. Collected information is stored in the Cross-Beta DB, and in CSV format.

Thus, for each entry we have the following information: its source and identity (Cross-Beta DB ID, PDB or AmyPro ID, protein name, species, etc.), the publication justifying its presence in the database (title, authors name, link/DOI/pmid, etc…), its sequence (AR sequence, predicted EAR, disordered region, mutation, AR containing protein sequence and ID, amino acid composition, etc.) and information about the conditions of amyloid formation (extraction method, observed concentration/pH/temperature, buffer, the method used (e.g. Congo-red birefringence, Electron Microscopy, ThT, etc…), number of protofibrils in the fibril and finally if the amyloid is known to be disease-related or functional). The aim is to allow the user to have all important information about a specific amyloid-forming region and generate features amenable for use in machine learning.

More details about the procedure of data integration in the database available in the supplementary data Figure S1.

### C. Cross-Beta DB characteristics

The Cross-Beta DB is currently composed of 116 non-redundant entries. They correspond to 44 different proteins from 14 different species (mainly *Homo sapiens*, with 99 entries) (Figure 2A, 2B). The majority of the entries are labeled as disease-related amyloids with some being functional or known as both disease-related and functional (Figure 2C). The amino acid composition analysis shows noticeable differences between sequences from the Cross-beta DB and globally from the Uniprot database (Figure 2D). Our analysis showed that the majority of the ARs have a length between 15 (the minimum required) 139 amino acids (DB00018, TAR DNA-binding protein 43), with an average of 56 amino acids (Figure 2E). The length distribution shows that typically ARs are about 5 times shorter than the average length of proteins (362 residues) from the reviewed Uniprot dataset.

**Figure 2:**
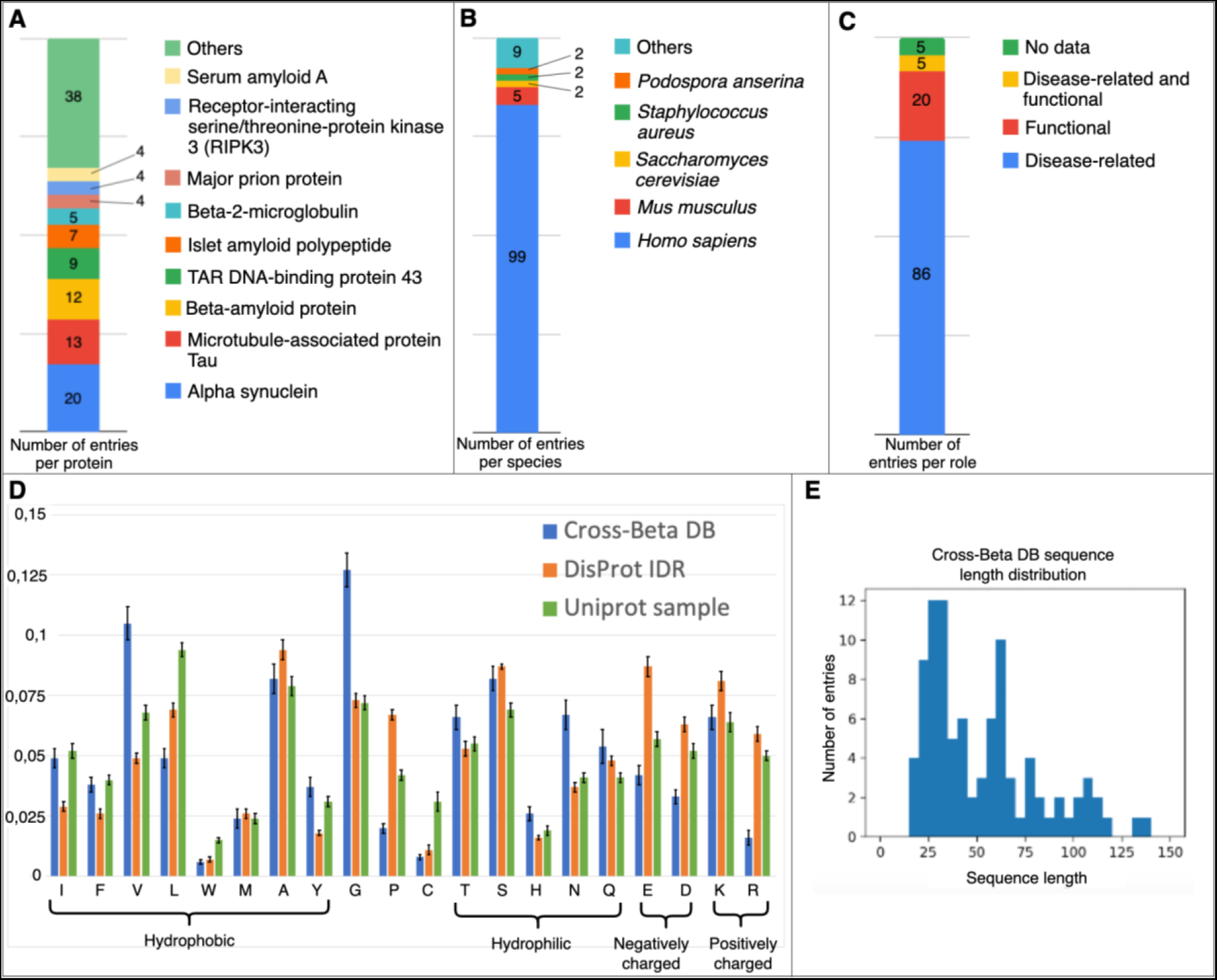
Composition details of the Cross-Beta DB. **(A)** Composition of the Cross-Beta DB in proteins where the “Others” value corresponds to proteins represented by 2 entries or less. **(B)** Composition of the Cross-Beta DB in species. The 14 different species are present but only 4 are represented by more than 1 entry. The other species are grouped in “*Others*”. **(C)** Composition of the Cross-Beta DB in protein roles. The “No data” value corresponds to entries where the pathogenicity is not explicitly proved. **(D)** Comparison of amino acid composition among the amyloid-forming regions of the Cross-Beta DB (Blue), the Intrinsically Disordered Regions of our negative set (see the next section) from DisProt (orange) and a sample of 150 random sequences of a length between 15 and 200 residues from UniProt (green) **(E)** Cross-Beta DB amyloid-forming region length distribution by number of entries. Values are grouped in buckets of 5 amino acids.

The Cross-Beta DB is available via a web interface at https://crossbetadb.crbm.cnrs.fr/index.html. It allows users to access and download the database data, as well as to filter the entries by categorical (e.g. “Role”) and numeric (e.g. “Sequence length”) values. The interface also features options for visualization of amyloid structures in the PDB and for analysis of amino acid composition.

### D. Construction of benchmarking datasets for amyloidogenicity predictors and training of machine learning models

To create a non-redundant positive dataset, we clustered sequences in the Cross-beta DB and randomly chose cluster representatives. The clustering was performed using CD-Hit (Fu et al., 2012) with an identity threshold of 0.70, and resulted in 53 clusters of significantly different amyloids. A non-redundant negative dataset was constructed from the DisProt database (Quaglia et al. 2022) using the same procedure. We selected 287 entries from 240 proteins, which resulted in 268 clusters each containing a representative (non-amyloid) Intrinsically Disordered Region (IDR) of length between 20 and 150 amino acids. This negative set was enriched with the same type of information as the positive set. These sets were used as is for the training of our machine learning models. Finally, we built 10 groups from the positive and negative sets, each composed of 10 randomly selected positive and 10 randomly selected negative sequences (see supplementary data Table S1) and used them for the benchmark.

### E. Developing a predictor based on machine learning by using Cross-Beta DB

#### Feature creation and selection

For training amyloidogenicity predictors on the Cross-Beta DB, we considered several features calculated using the amino acid sequence of each entry, which were then reduced via feature selection in order to retain only the features with the most impact on model prediction accuracy and to remove those with neutral or negative effects, while also reducing the train and inference time of the algorithm. The considered features fell into two broad categories (Figure 3): features that were calculated directly from the amino acid sequence and features that were calculated from reduced representations of the amino acid sequence obtained by categorizing sets of amino acids into groups.

**Figure 3:**
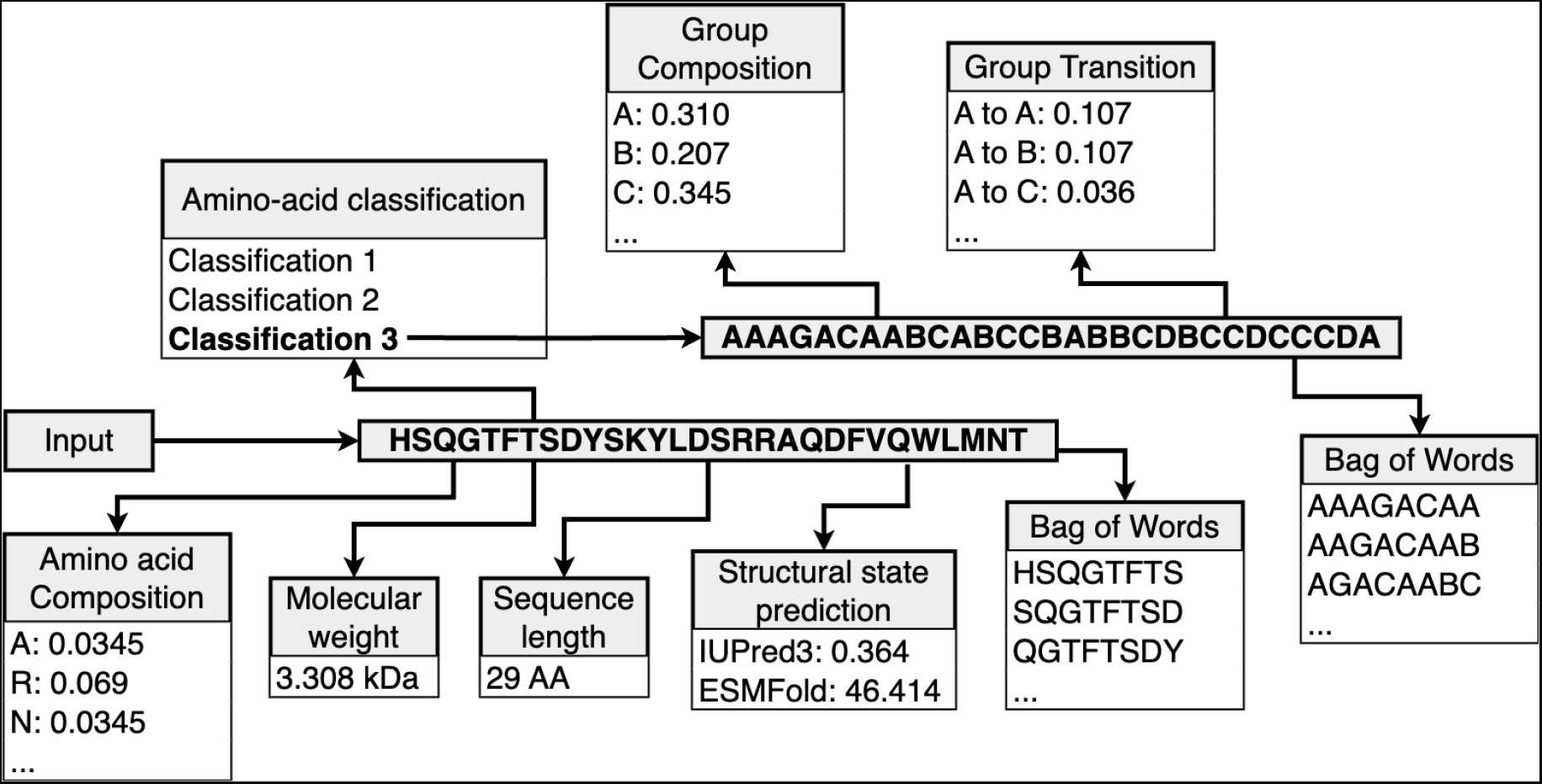
Process of feature creation using an amino acid sequence as input

**Figure 3:**
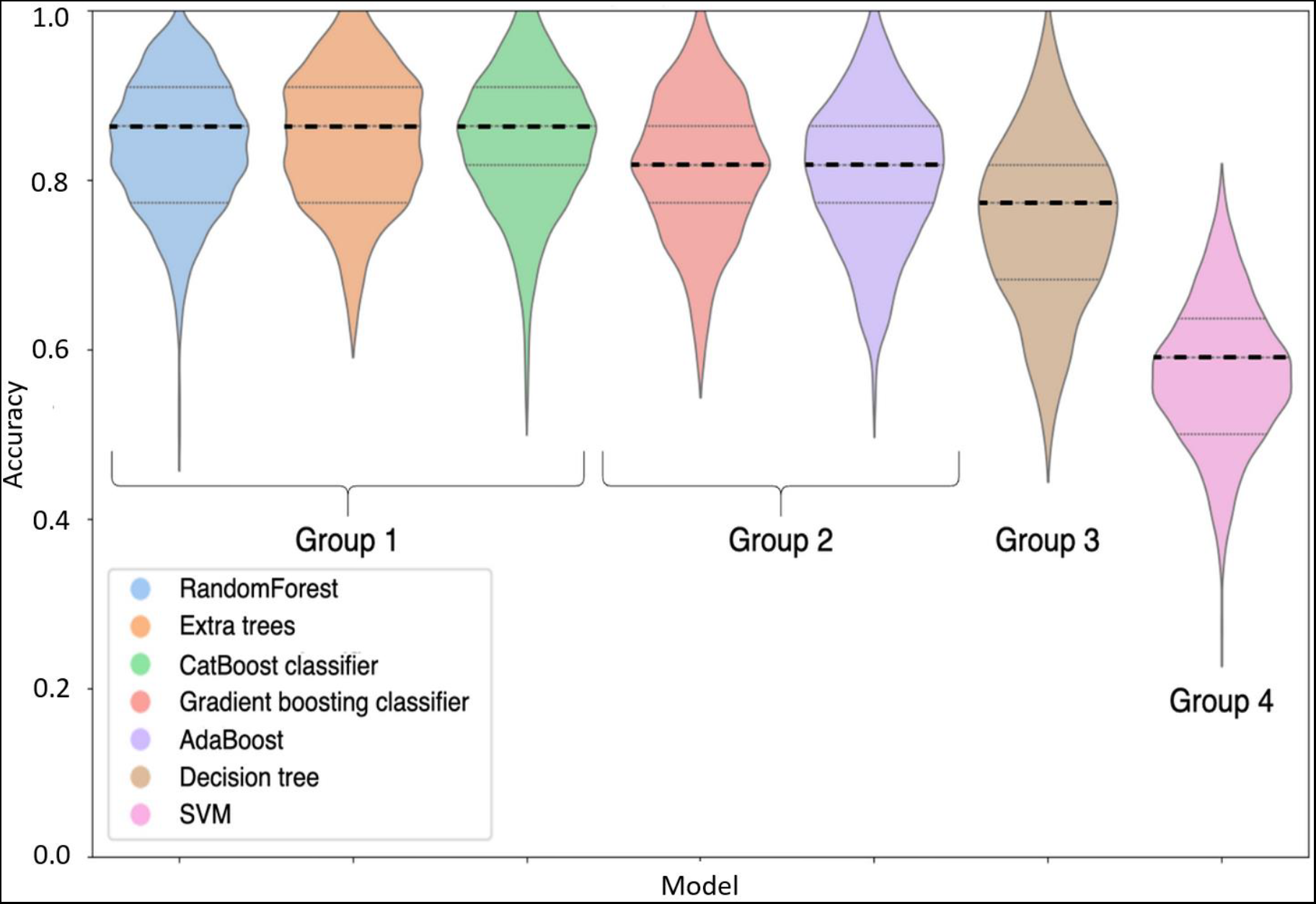
Prediction accuracy for the 7 selected classifiers: Random Forest, Extra trees, CatBoost classifier, Gradient boosting classifier, Adaboost, Decision tree, and Support Vector Machine (SVM). Four significantly different groups can be isolated: group 1 (RandomForest, Extra Trees and CatBoost), group 2 (Gradient boosting classifier, Adaboost), group 3 (Decision tree) and group 4 SVM. All the groups are significantly (p-value < 0.0001) different from each other. All grouped models share no significant difference (p-value > 0.05) with models of the same group (see supplementary data S3). Accuracy distribution comparison was realized using the Wilcoxon-Mann-Whitney test.

The directly calculated features were as follows: sequence length, molecular weight, amino acid composition (fraction of each amino acid present), pairwise amino-acid transition frequencies (fraction of each amino acid pair present), potential to form stable structure (pLDDT from ESMFold structure predictions and disorder probability from IUPred3), and presence of k-mers (k=8) generated using the Bag of Words technique (BoW) (Zhang et al., 2010).

In addition to the directly calculated features, we considered features based on grouped representations of each sequence. To do this we considered three classification methods (Table S2). Classification 1 was taken from Chothia and Finkelstein amino acid classification (Chothia & Finkelstein, 1990) based on their preference to be in the different types of the secondary structure. Classification 2 represented a typical subdivision of amino acids based on their physico-chemical properties. Classification 3 was suggested based on known amyloidogenicity properties of amino acid residues (Ahmed et al., 2015). For example, in accordance with Classification 1 an amino acid sequence “ACDEFGHI” was represented as “BCAACBBC” (Table S2). Using these amino acid groups, we calculated features analogous to those calculated on the raw sequence, namely: group composition, group transitions, and bag-of-words features. We found that Classification 3 provided the best accuracy in all cases, so we used only this Classification for further model development.

Features were selected using the built-in feature selection methods of the Random Forest Scikit-learn model (Pedregosa et al., 2011): once the model is trained and tested, the algorithm selects a feature and replaces its value by random ones. The accuracy before and after the change are compared and the change is stored. This process considers one feature at a time and gives, at the end, the accuracy change for each of the tested features. The final set of chosen features is shown in Table 1.

**Table 1:** Set of features selected by the feature importance algorithm.

| Feature type | Features |
| --- | --- |
| Amino acid composition | amino acids N, D, C, Q, I, L, M, F, T, W, and Y |
| Amino acid group composition | groups A, B, C, and G |
| Amino acid group transitions | $A \rightarrow A$ and $A \rightarrow C$<br>$B \rightarrow A$, $B \rightarrow B$, $B \rightarrow D$, and $B \rightarrow G$<br>$C \rightarrow A$, $C \rightarrow B$, $C \rightarrow C$, $C \rightarrow P$ and, $C \rightarrow G$<br>$D \rightarrow B$, $D \rightarrow C$ and, $D \rightarrow G$<br>$P \rightarrow B$, $P \rightarrow C$, $P \rightarrow D$, $P \rightarrow P$ and, $P \rightarrow G$<br>$G \rightarrow A$, $G \rightarrow B$ and, $G \rightarrow G$ |
| Structural state prediction | IUPred score |

#### Model selection

We trained and tested five Ensemble classifiers as well as Decision tree and Support Vector Machine (SVM) models on the Cross-Beta DB (Figure 3). The models have been compared by using the default values of provided hyper-parameters. To evaluate their performance we used two metrics: Accuracy = (TP+TN) / (TP+TN+FP+FN) and F1 score = 2 *TP/(2TP+FN+FP)) where TP is the number of true positives, TN - true negatives, FP - false positives, and FN - false negatives. Both metrics led us to the same conclusion. On Figure 3, to demonstrate the results of the tests, we use the accuracy. In addition to the accuracy comparison, we evaluated the difference between the number of TP and FP (Figure S2) for each model depending on the chosen threshold. Statistical comparisons have been made on the accuracy using Wilcoxon-Mann-Whitney statistical non-parametric tests. Our results suggested that three models (Random Forest, Extra trees, CatBoost classifier) perform equally well and outperform the others (Figure 3 and Figure S2, Table S3, Figure S3). They have been chosen for the next step of hyper-parameter optimization.

We then optimized the hyper-parameters of each model using a grid search. After conducting the comparison of the optimized models, we chose the Random Forest model as the most effective predictor and named it Cross-Beta RF.

#### Cross-Beta RF predictor usage

The Cross-Beta RF predictor takes as input an amino acid sequence of length 15 AAs or more, which is in turn used to generate all the features. The output consists of: average score of the query prediction, positions of predicted ARs having scores of higher than the chosen threshold (default threshold is 0.5) and the minimal length of 15 AA, and the value of the score for each amino acid of the sequence. This amino acid score is computed using a window of a chosen length (minimum 15 residues and can’t be longer than the input sequence). While the window slides from N-term to C-term of the sequence, the prediction is run for each new sequence fragment within the window. Each amino acid has a different score for each fragment sequence it has been part of, the average of these scores give the amino acid final score. The output also consists of a graph representing the chosen threshold with all the amino acids scores of the query sequence. The Cross-Beta RF predictor is available via a web page: https://bioinfo.crbm.cnrs.fr/index.php?route=tools&tool=35.

### F. Benchmark of existing amyloid predictors

Using our benchmark sets we compared performance of our Cross-beta DB predictors with several other popular and available methods, which are based on different approaches: Tango, a knowledge based amyloid predictor, that uses secondary structure prediction (Fernandez-Escamilla et al., 2004), ArchCandy 2.0 a knowledge-based method that leverages physico-chemical properties of amino acids to predict amyloid-forming β-arches (Ahmed et al., 2015), Pasta 2.0 a knowledge based amyloid predictor reinforced with machine learning generated features (Walsh et al., 2014), and AmyloGram, a machine learning amyloid predictor based on a Random Forest model (Burdukiewicz et al., 2017). Our benchmark was performed, using the 10 generated groups each containing 10 different positive (ARs from Cross-Beta DB) and 10 different negative (IDRs from DisProt) randomly selected entries from our benchmark set (see section D).

All predictors were used with their default settings. ArchCandy 2.0 was used with a threshold of 0.4. Tango with its default settings was used through the TAPASS pipeline (Falgarone et al., 2022). Pasta 2.0 was used with the default setting “Region (85% spec)”, which gave the best results between 3 suggested settings. AmyloGram was used online with its default settings (Cut off = 0.5). Cross-Beta RF has been used with default settings provided by the Python library Scikit-learn.

To evaluate predictors, in addition to the F1 score and accuracy, we used recall metrics TP/(TP+FN) and precision metrics TP/(TP+FP). The comparison of the predictors showed that Cross-beta RF predictor, with accuracy 0.837 and F1 score 0.845, demonstrated the best performance (Table 2). The other programs ranked in order of better performance exhibit the following results: ArchCandy 2.0 obtains an accuracy of 0.720 and an F1 score of 0.759 while Pasta 2.0, Tango, and AmyloGram have an accuracy of around 0.5 - 0.6 (Table 2).

**Table 2:** Results summary for all tested amyloid prediction methods. Among Pasta2.0 default settings, it is the settings 3 that perform the best (in accuracy and F1 score) (Table S1 and Table S4). The Random Forest corresponds to the machine-learning model we are developing for the prediction of amyloids.

| Prediction method | Accuracy | Recall | Precision | F1 score |
| --- | --- | --- | --- | --- |
| Cross-Beta RF | 0.837 | 0.859 | 0.831 | 0.845 |
| ArchCandy2.0 | 0.720 | 0.870 | 0.673 | 0.759 |
| Pasta2.0 (Settings 3) | 0.520 | 0.880 | 0.513 | 0.648 |
| AmyloGram | 0.585 | 0.750 | 0.567 | 0.646 |
| Tango | 0.550 | 0.310 | 0.590 | 0.406 |

Thus, the newly developed Cross-beta RF predictor shows the best results in terms of accuracy and F1 score with the data from the Cross-beta DB.

## Conclusion

Proteins have the capability to form aggregates, which can manifest in diverse forms, ranging from amorphous to fibrous structures (Fink, 1998). Fibrillar aggregates, in turn, display various structures, including fibrils formed by globular structures, alpha-helical fibrils, and, ultimately, beta-structural fibrous formations. Among these, beta-structural aggregates with cross-β structures, known as amyloids, are the most abundant. Finally, the cross-β structures are not uniform and can be subdivided into two groups. One group consists of amyloids formed naturally in vivo, often associated with human diseases. These amyloids require a certain minimal length of the amyloid-forming motif, around 15 residues. The other group includes cross-b amyloid fibrils formed by short peptides typically ranging between 4-10 residues. These fibrils are formed at higher concentrations in vitro and often under specific non-physiological conditions.

Without a doubt, naturally occurring amyloids, especially those associated with diseases, are of paramount importance to biologists and medical professionals. To understand their structural principles and predict their existence in analyzed proteins, a special database of naturally occurring amyloids is needed. However, until now, existing databases have represented a mixture of different types of protein aggregates, partially explaining the poor performance of prediction methods developed on this data.

In this study, we surveyed existing databases, scrutinized the data, and constructed a Cross-beta DB specifically containing information about amyloidogenic motifs capable of forming amyloids under natural conditions. The Cross-beta DB, featuring denoised data, has not only paved the way for the development of more effective machine learning tools for predicting naturally occurring amyloids but has also provided a platform for the comparative analysis of predictors.

As a result, by using the Cross-beta DB, we developed a machine learning based tool, Cross-Beta RF, capable of predicting amyloidogenicity of a given amino acid sequence. The benchmark demonstrated the superior performance of the Cross-beta RF predictor over other existing methods. In the future, the Cross-beta DB, with its criteria for entry selection, can be maintained, reproduced, and expanded. This database is poised to facilitate further progress in the development of improved methods for predicting the most crucial type of protein aggregates-naturally occurring amyloid fibrils.

## Acknowledgments

This work was supported in part by a CIFRE PhD fellowship through the “*Association Nationale de la Recherche Technique*” (ANRT) program, in collaboration between PROTERA and “*Centre National de la Recherche Scientifique*” (CNRS) to V.G. and by EU COST Action ML4NGP CA21160 to V.G and A.V.K. The authors thank Dr Layla Hirsh for critical reading of the manuscript, comments and for assistance with the implementation of Cross-Beta RF predictor web-interface. We also thank Mrs Stefany Neciosup Vera for assistance with statistical analysis of the data.

## Data availability

Cross-beta DB is available online: https://crossbetadb.crbm.cnrs.fr/index.html and Cross-beta RF prediction is available online: https://bioinfo.crbm.cnrs.fr/index.php?route=tools&tool=35

## Notes

https://crossbetadb.crbm.cnrs.fr/index.html

https://bioinfo.crbm.cnrs.fr/index.php?route=tools&tool=35

### Competing Interest Statement

The authors have declared no competing interest.

