## Supplemental Files for "Developing machine-learning-based amyloid predictors with Cross-Beta DB"

#### Development of machine learning based amyloid predictor by using Cross-Beta DB database.

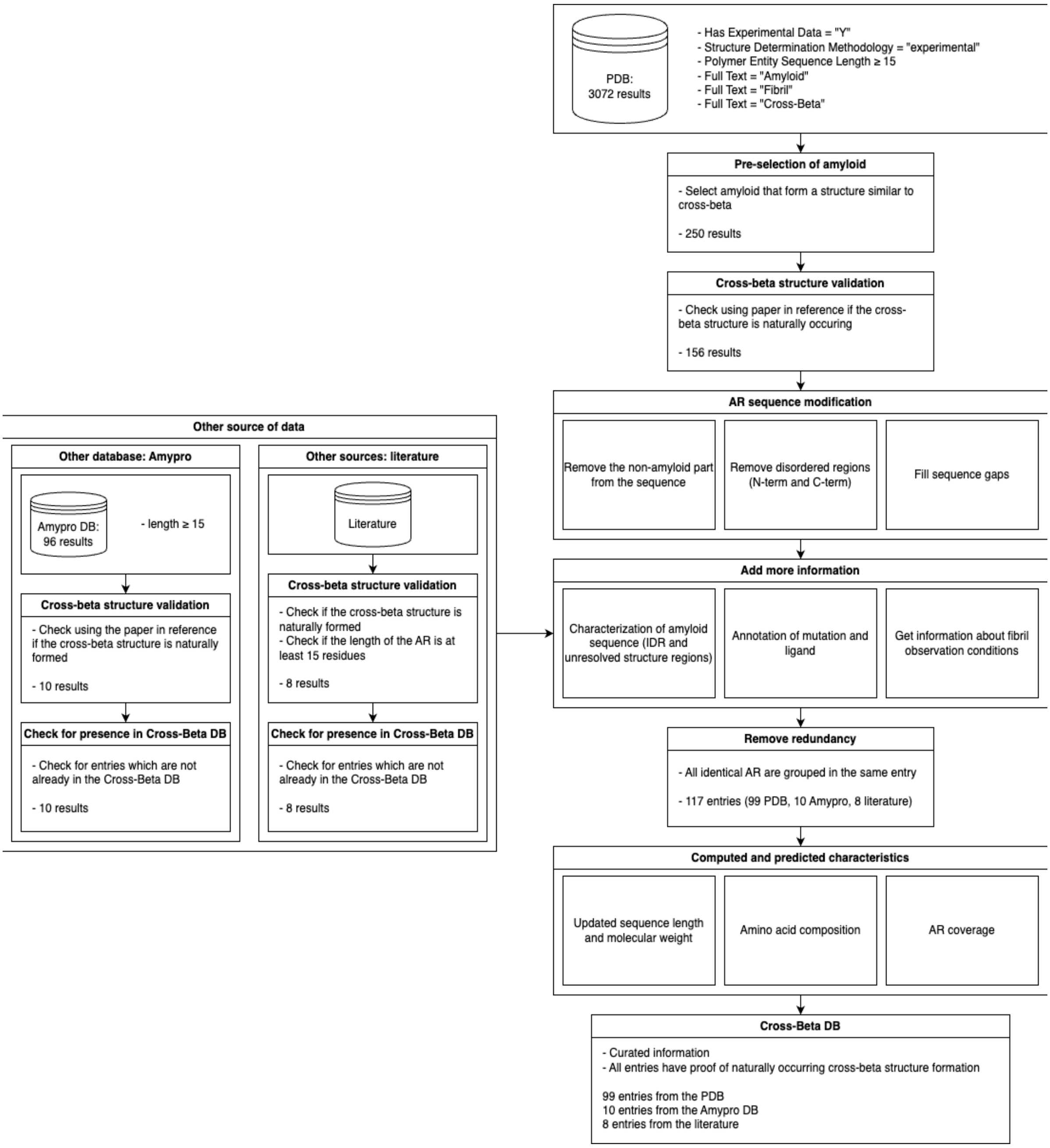

Figure S1: Data cleaning pipeline

### Table S1: Benchmark test groups:

| Set 1 | Positive | Negative |
|---|---|---|
| 1 | ANFLVHSSNNFGAILSSTNVGSNTYX | TESLVLSPAPAKPKRVKASRR |
| 2 | RPLVNIYNCSGVQVGDNNYLTMQQTTALP | ATRLTKKQLAQAIQNTLPNPPRRKRRAKRRAAQVPKPTQAGVSM<br>APIAQGTMVKLRPPMLRSSM |
| 3 | MSAEGYQYRALYDYKKEREEDIDLHLGDILTVNKGSLVALGFSDG<br>QEAKPEEIGWLNGYNETTGERGDFPGTYVEYI | MSQKPLSDDEKFLFVDKNFVNNPLAQADWSAKK |
| 4 | KVQIINKKLDLSNVQSKCGSKDNIKHVPGGGSVQIVYKPVDLSKVT<br>SKCGSLGNIHHKPGGGQVEVKSEKLDFKDRVQSKIGSLDNITHVP<br>GGGNKKIETHKLTFRE | RVKFSRSADAPAYQQGQNQLYNELNLGRREEYDVLDKRRGRDP<br>EMGGKPQRRKNPQEGLYNELQKDKMAEAYSEIGMKGERRRGKG<br>HDGLYQGLSTATKDTYDALHMQALPPR |
| 5 | GSVQIVYKPVDLSKVTSKCGSLGNIHHKPGGGQVEVKSEKLDFKD<br>RVQSKIGSLDNITHVPGGGNKKIETHKLTFRE | MKVLILACLVALALARELEELNVPG |
| 6 | KVQIINKKLDLSNVQSKCGSKDNIKHVPGGGSVQIVYKPVDLSKVT<br>SK | DGLVGMAIVGGMALGVAGLAGLIGLAVSKSKS |
| 7 | YVGSKTKEGVVHGVATVAEKTKEQVTNVGGAVVTGVTAVAQKTV<br>EGAGSIAAATGFVKK | MPREFKSFGSTEKSLLSKGHGEPSYSEI |
| 8 | QKLVFFAEDVGSNKGAIIGLMVGGVVIA | LTAATHWASMDPAVVHPELNGAAYSRYPPGVVSVAPTGIPAAVE<br>GIVPSAMSLSHGLPPVAHPPHAPSPGQTVKPEADRDHTTDQL |
| 9 | MNTLFNLFFDFITGILKNIGNIAA | MATANSIIVLDDDDEDEAAAQPGPSHPLPNAASPGAEAPSSSEPH<br>GARGSSSSG |
| 10 | GSTLGTLGGAAVGGVIG | TTPPSVYPLAPGSAAQTNSMVTLGCLVKGYFPEPVTVTWNSGSL<br>SSGVHTFPAVLQSDLYTLSSSVTVPSSTWPSETVTCNVAHPASST<br>KVDKKIVPR |

| Set 2 | Positive | Negative |
|---|---|---|
| 1 | SYSGYSQSTDTSGYGQSSYSSYGQSQNTGYGTQSTPQGYGSTG<br>GYGSSQSSQSSYGQQSSY | MARYRRSRTRSRSPRSRRRRRRSGRRRSPRRRRRYGSARRSR<br>RSVGGRRRRYGSRRRRRRY |
| 2 | MNFGAFSINPAMMAAAQAALQSSWGMMGMLASQQNQ | KADMNTFPNFTFEDPKFEVVEKPQS |
| 3 | GFGNSRGGGAGLGNNQGSNMGGGMNFGEFSIN | PQTPSRPASEQPPAQPRLRIAEQDPNWETF |
| 4 | GSNFGGGGSYNDFGNYNNQSSNFGPMKGGN | QRETSAVVKTVTPAAPRKMAVADSEENWETF |
| 5 | HSQGTFTSDYSKYLDSRRAQDFVQWLMNT | MPKSKELVSSSSSGSDSDSEVDKKLKRKKQVAPEKPVKKQKTGE<br>TSRALSSSKQSSSSRD |
| 6 | VFNNCSEVQIGNYNSLVAPPR | AKDLLKADDIKKALDAVKAEGSFNHKKFFALVGLKAMSANDVKKV<br>FKAIDADASGFIEEEELKFVLKSFAADGRDLTDAETKAFLKAADKD<br>GDGKIGIDEFETLVHEA |
| 7 | MSAEGYQYRALYDYKKEREEDIDLHLGDILTVNKGSLVALGFSDG<br>QEAKPEEIGWLNGYNETTGERGDFPGTYVEYI | YRKDKRRQEPLRQPSPQRGAGAPELGAAPEEELAALQLGPTHHE<br>CEAGPPHDTLRLTALPDYTLTLRRSPDDIPLMTPNTITMIPNSLVGL<br>QTLHPYNTFAAGFNSTGLPHSHSTTRV |

| | | |
|---|---|---|
| 8 | QKLVFFAENVGSNKGAIIGLMVGGVV | MLFIKPADLREIVTFPLFSDLVQCGFPSPAADYVEQRIDLNQLLIQHPSATYFVKASGDSMIDGGISDGDLLIVDSAITASHGDIVIAAVDGEFTVKKLQLRPTVQLIPMNSAYSPITISSEDTLDVFGVVIHVVKAMR |
| 9 | MGIIAGIIKVIKSLIEQFTGK | MSLIPSFFSGRRSNVFDPFSLDVWDPLKDFPFSNSSPSASFPRENPAFVSTRV |
| 10 | NGNGASQYFGNSMTTGNMSPQMALIQGSFNKP | GGDSDNEKGGKSSKKRKTEPSPSKKANTSGSGFKSKEYISDDDSTSSDDEKDNEPAKKKSKPPSDGDAKKKKAKSESEPEESEEDSNASDEDEEDEASD |

| Set 3 | Positive | Negative |
|---|---|---|
| 1 | ATAVSEWTEYKTADGKTFYYNNRTLE | VDSVYRTRSLGVAAEGIPDQYADGEAARVWQLYIGDTRSR |
| 2 | GFGNSRGGGAGLGNNQGSNMGGGMNFGEFSIN | MSESSSKSSQPLASKQEKDGTEKRGRGRPRKQPPVSPGTALVGSQKEPSEVPTPKRPRGRPKGSKNKGAAKTRKTTTTPGRKPRGRPKKLEKEEEEGISQESSEEEQ |
| 3 | GYGNQGGGYGGGYDNYGGGNYGSGNYNDFGNYNQQPSNYGPMKSGNFGGSRNMGGPY | MCKGLAGLPASCLRSAKDMKHRLGFLLQKSDSCEHSSSHSKKDKVVTCQR |
| 4 | THNQWNKPSKPKTNMKHMAGAAAAGAVVGGLGGYMLGSAMSRPMMHFGNDWEDRYYRENMNRYPNQVYYRPVDQYNNQNNFVHDCVNITIKQHTVTTTTKGENFTETDIKIMERVVEQMCTTQYQKESQAYYD | MEEPQSDPSVEPPLSQETFSDLWKLLPENNVLSPLPSQAMDDLMLSPDDIEQWFTEDPGPDEAPRMPEAAPPV |
| 5 | SNQNNFVHDCVNITIKQHTVTTTTKGENFTETDVKMMERVVEQMCITQYERESQAYYQRG | MAEAGAGLSETVTETTVTVTTEPENRSLTIKLRKRKPEKKVEWTSDTVDNEHMGRRSSKCCCIYEKPRAFGESSTESDEEEEEGCGHTHCVRGHRKGRRRATLGPTPTTPPQPPDPSQPPPGPMQH |
| 6 | GSVQIVYKPVDLSKVTSKCGSLGNIHHKPGGGQVEVKSEKLDFKDRVQSKIGSLDNITHVPGGGNKKIETHKLTFRE | MKVLILACLVALALARELEELNVPG |
| 7 | GGKVQIVYKPVDLSKVTSKCGSLGNIHH | MKAKRSHQAIIMSTSLRVSPSIHGYHFDTASRKKAVGNIFENTDQESLERLFRNSGDKKAEERAKIIFAIDQDVEEKTRALMALKKRTKDKLFQFLKLRKYSIKVH |
| 8 | RSVQTIVFQPQLASRTPTGQS | MKKRNRLKKNEDFQKVFKHGTSVANRQFVLYTLDQPENDELRVGLSVSKKIGNAVMRNRIKRLIRQAFLEEKERLKEKDYIIIARKPASQLTYEETKKSLQHLFRKSSLYKKSSSK |
| 9 | NGNGASQYFGNSMTTGNMSPQMALIQGSFNKP | PRKQVSGPERTIPITREEKPAVTAAPKK |
| 10 | INDIIFNTNLANNLSNYN | WNRAKLSLAQKKDRVAQKKASFLRAQEKADS |

| Set 4 | Positive | Negative |
|---|---|---|
| 1 | GQRVTISCSGRSSNIGRNLVKWYQQFPGTAPKLLIYSNDQRPSGVPDRFSGSKSGTSASLAVSGLQSEDEADYYCAAWDATLNAWVFGGGT | MSTEASVSYAALILADAEQEITSEKLLAITKAAGANVDQVWADVFAKAVEGKNLKELLFSFAAAAPASGAAAGSASGAAAGGEAAAEEAAEEEAAEESDDDMGFGLFD |
| 2 | QLHQQQHQQQHQQHQQHQQQQQLHQHQQQLS | PKRKVSSAEGAAKEEPKRRSARLSAKPAPAKVETKPKKAAGKDKSSDKKVQTKGKRGAKGKQAEVANQETKEDLPAENGETKNEESPASDEAEEKEAKS |
| 3 | SYGSSSQSSSYGQPQSGSYSQQPSYGGQQQSYGQQQSYN | MEEDYSWAEEEDEGEAEGEEEEEEEEDQESPPKAVKRPAATKKAGQAKKKKLDKEDESSEEDSPTKKGKGAGRGRKPAAKK |
| 4 | MNFGAFSINPAMMAAAQAALQSSWGMMGMLASQQNQ | MKVLILACLVALALARELEELNVPG |

| | | |
|---|---|---|
| 5 | GSNFGGGGSYNDFGNYNNQSSNFGPMKGGN | MNLEPPKAEIRSATRVMGGPVTPRKGPPKFKQRQTRQFKSKPPKKGVQGFGDDIPGMEGLGTDITVICPWEAFNHLELHELAQYGII |
| 6 | VFNNCSEVQIGNYNSLVAPPR | EGRQYGRGRSGGQVRDEYRQDYDAGRGGYGKLAQNQ |
| 7 | THNQWNKPSKPKTNMKHMAGAAAAGAVVGGLGGYMLGSAMSRPMMHFGNDWEDRYYRENMNRYPNQVYYRPVDQYNNQNNFVHDCVNITIKQHTVTTTTKGENFTETDIKIMERVVEQMCTTQYQKESQAYYD | LTAATHWASMDPAVVHPELNGAAYSRYPPGVVSVAPTGIPAAVEGIVPSAMSLSHGLPPVAHPPHAPSPGQTVKPEADRDHTTDQL |
| 8 | YVGSKTKEGVVHGVATVAEKTKEQVTNVGGAVVTGVTAVAQKTVEGAGSIAAATGFVKK | VKSDGNILDDLNEATKKASDFVTDKTKEALADGEKTKDYIVEKTIEANETATEEAKKALDYVTEKGKEAGNKAAEFVEGKAEEAKNATKS |
| 9 | QKLVFFAENVGSNKGAIIGLMVGGVV | MATANSIIVLDDDDEDEAAAQPGPSHPLPNAASPGAEAPSSSEPHGARGSSSSG |
| 10 | HYEYKSYNAGHNVESVVENKLVDASDLTLGVDIL | GGRQGGGAPAGGNIGGGQPQGGWGQPQQPQGGNQFSGGAQSRPQQSAPAAPSNEPPMDFDDDIPF |

| Set 5 | Positive | Negative |
|---|---|---|
| 1 | SFFSFLGEAFDGARDMWRAYSDMREANYIGSDKYFHARGNYDAAKRGPGGVWAA | MSATAATAPPAAPAGEGGPPAPPPNLTSNRRLQQTQAQVDEVVDIMRVNVDKVLERDQKLSELDDRADALQAGASQFETSAAKLKRKYWWKNLKMM |
| 2 | PLMVKVLDAVRGSPAINVAMHVFRKAADDTWEPFASGKTSESGELHGLTTEEEFVEGIYKVEIDTKSYWKALGISPFHEHAEVVFTANDSGPRRYTIAALLSPYSYSTTAVVT | MRLRKYNKSLGWLSLFAGTVLLSGCN |
| 3 | ATAVSEWTEYKTADGKTFYYNNRTLE | MSTQQQARALMMRHHQFIKNRQQSMLSRAAAEIGVEAEKDFWTTVQGKPQSSFRTTYDRSNASLS |
| 4 | GFGNSRGGGAGLGNNQGSNMGGGMNFGEFSIN | WAAEEKRIPSSGDLSESDDWSEEPKQ |
| 5 | ANFLVHSSNNFGAILSSTNVGSNTYX | MSGGDGRGHNTGAHSTSGNINGGPTGIGVSGGASDGSGWSSENNPWGGGSGSGIHWGGGSGRGNGGGNGNSGGGSGTGGNLSA |
| 6 | GYGNQGGGYGGGYDNYGGGNYGSGNYNDFGNYNQQPSNYGPMKSGNFGGSRNMGGPY | MSLEKAHTSVKKMTFGENRDLERVVTAPVSSG |
| 7 | KVQIINKKLDLSNVQSKCGSKDNIKHVPGGGSVQIVYKPVDLSKVTSKCGSLGNIHHKPGGGQVEVKSEKLDFKDRVQSKIGSLDNITHVPGGGNKKIETHKLTFRE | MDRSLGWQGNSVPEDRTEAGIKRFLEDTTDDGELSKFVKDFSGNASCHPPEAKTWASRPQVPEPRPQAPDLYDDDLEFRPPSRPQSSDNQQYFCAPAPLSPSARPRSPWGKLDPYDSSE |
| 8 | KTNMKHMAGAAAAGAVVGGLGGYMLGSAMSRPIIHFGSDY | NDWFSKLASSAFSGLFGALLA |
| 9 | GVVAAAEKTKQGVAEAAGKTKEGVLYVGSKTKEGVVHGVATVAEKTKEQVTNVGGAVVTGVTAVAQKTVEGAGSIAAATGFVKK | MSATAATVPPAAPAGEGGPPAPPPNLTSNRRLQQT |

| Set 6 | Positive | Negative |
|---|---|---|
| 1 | NFMLTQPHSVSESPGKTLTISCTGSSASIASHYVQWYQQRPGGAPTTLIYENDQRPSEVPDRFSGSIDSSSNSASLTISGLKTEDEADYYCQSYDGNNHWVFGGG | PKRKSATKGDEPARRSARLSARPVPKPAAKPKKAAAPKKAVKGKKAAENGDAKAEAKVQAAGDGAGNAK |
| 2 | GFGNSRGGGAGLGNNQGSNMGGGMNFGEFSIN | HIKKPLNAFMLYMKEMRANVVAECTLKESAAINQILGRRWHALSREEQAKYYELARKERQLHMQLYPGWSARDNYGKKKKRKREKLQESTSGTGPRMTAAYI |

| | | |
|---|---|---|
| 3 | GYGNQGGGYGGGYDNYGGGNYGSGNYNDFGNYNQQPSNYGPMKSGNFGGSRNMGGPY | MEEPQSDPSVEPPLSQETFSDLWKLLPENNVLSPLPSQAMDDLMLSPDDIEQWFTEDPGPDEAPRMPEAAPPV |
| 4 | FLNCYVSGFHPSDIEVDLLKNGERIEKVEHSDLSFSKDWSFYLLYYTEFTPTEKDEYACRVNHV | MDRTTTRLYAGLKKRLAMASANKA |
| 5 | THNQWNKPSKPKTNMKHMAGAAAAGAVVGGLGGYMLGSAMSRPMMHFGNDWEDRYYRENMNRYPNQVYYRPVDQYNNQNNFVHDCVNITIKQHTVTTTTKGENFTETDIKIMERVVEQMCTTQYQKESQAYYD | MEEDYSWAEEEDEGEAEGEEEEEEEEEDQESPPKAVKRPAATKKAGQAKKKKLDKEDESSEEDSPTKKGKGAGRGRKPAAKK |
| 6 | KVQIINKKLDLSNVQSKCGSKDNIKHVPGGGSVQIVYKPVDLSKVTSKCGSLGNIHHKPGGGQVEVKSEKLDFKDRVQSKIGSLDNITHVPGGGNKKIETHKLTFRE | RYLHTALEGMANPEDPECESEGWLLEKSVPETWKAFLESVKKLGKGNQVEAEGEDAGQAPAAG |
| 7 | MNTLFNLFFDFITGILKNIGNIAA | MPSLMSFGSCQWIDQGRFSRSLYRNFKTFKLHEMHGLC |
| 8 | DQLETQTRELETAYSNLLRD | ISLAGKTNFFEKRVSDYQKAGVMSKSTKQEAGAFTFNEDF |
| 9 | HYEYKSYNAGHNVESVVENKLVDASDLTLGVDIL | YNAVNPFEFMEDVATAGKTTFFEKKVSDYQKASDMSKSATPSKEINFDDDF |
| 10 | NGNGASQYFGNSMTTGNMSPQMALIQGSFNKP | SLRRSSCFGGRMDRIGAQSGLGCNSFRY |

| Set 7 | Positive | Negative |
|---|---|---|
| 1 | NFMLTQPHSVSESPGKTLTISCTGSSASIASHYVQWYQQRPGGAPTTLIYENDQRPSEVPDRFSGSIDSSSNSASLTISGLKTEDEADYYCQSYDGNNHWVFGGG | MTDVETTYADFIASGRTGRRNAIHDILVSSASGNSNELALKLAGLDINKTEGEEDAQRSSTEQSGEAQGEAAKSE |
| 2 | SFFSFLGEAFDGARDMWRAYSDMREANYIGSDKYFHARGNYDAAKRGPGGVWAA | NPSSCGAEKQKGAKSSADCTSLVPQCA |
| 3 | PLMVKVLDAVRGSPAINVAMHVFRKAADDTWEPFASGKTSESGELHGLTTEEEFVEGIYKVEIDTKSYWKALGISPFHEHAEVVFTANDSGPRRYTIAALLSPYSYSTTAVVT | MPRSLKKGPFIDLHLLKKVEKAVESGDKKPLRTWSRRSTIFPNMIGLTIAVHNGRQHVPVFVTDEMVGHKLGEFAPTRTYRGHAADKKAKKK |
| 4 | GFGNSRGGGAGLGNNQGSNMGGGMNFGEFSIN | MPQLNGGGGDDLGANDELISFKDEGEQEEKSSENSSAERDLADVKSSLVNESETNQ |
| 5 | ANFLVHSSNNFGAILSSTNVGSNTYX | MSSQQNQNRQGEQQEQGYMEAAKEKVVNAWESTKETLSSTAQAAAEKTAEFRDSAGETIRDLTGQAQEKGQEFKERAGEKAEETKQRAGEKMDETKQRAGEMRENAGQKMEEYKQQGKGKAEELRDTAAEKLHQAGEKVKGRD |
| 6 | THNQWNKPSKPKTNMKHMAGAAAAGAVVGGLGGYMLGSAMSRPMMHFGNDWEDRYYRENMNRYPNQVYYRPVDQYNNQNNFVHDCVNITIKQHTVTTTTKGENFTETDIKIMERVVEQMCTTQYQKESQAYYD | EPVRKEPVFTLAELVNDITPENLHENIDWGEPKDKEVW |
| 7 | KVQIINKKLDLSNVQSKCGSKDNIKHVPGGGSVQIVYKPVDLSKVTSK | MAFSARFPLWLLLGVVLLASVSASFAHSGHSGGEAEDESEESRAQ |
| 8 | PLVNIYNCSGVQVGD | SDEIMDLLVQSVTKSSPRALAARERKRSRGNRKSLRRTLKSGLGDDLVQALGLSKGPGLEV |
| 9 | QKLVFFAENVGSNKGAIIGLMVGGVV | RLKIQVRKAAITSYEKSDGVYTGLSTRNQETYETLKHEKPPQ |
| 10 | MGIIAGIIKVIKSLIEQFTGK | GAGQSSPATGSQNQSGNTGSIINNYYMQQYQNSMDTQLG |

| Set 8 | Positive | Negative |
|---|---|---|
| 1 | GQRVTISCSGRSSNIGRNLVKWYQQFPGTAPKLLIYSNDQRPSGVPDRFSGSKSGTSASLAVSGLQSEDEADYYCAAWDATLNAWVFGGGT | MSSSTPFDPYALSEHDEERPQNVQSKSRTAELQAEIDDTVGIMRDNINKVAERGERLTSIEDKADNLAVSAQGFKRGANRVRKAMWYKDLKMK |
| 2 | SYSGYSQSTDTSGYGQSSYSSYGQSQNTGYGTQSTPQGYGSTGGYGSSQSSQSSYGQQSSY | TSHVQEEQIEVEETIEAAKAEEAKDEPPSEGEAEEEGKEKEEAEAEAEAEEEGAQEEEEAAEKEESEEAKEEEGGEGEQGEETKEAEEEEKKDEGAGEEQATKKKD |
| 3 | MNFGAFSINPAMMAAAQAALQSSWGMMGMLASQQNQ | MSTESALSYAALILADSEIEISSEKLLTLTNAANVPDENIWADIFAKALDGQNLKDLLVNFSAGAAAPAGVAGGVAGGEAGEAEAEKEEEEAKEESDDDMGFGLFD |
| 4 | GFGNSRGGGAGLGNNQGSNMGGGMNFGEFSIN | LNKWKSKGRRFKGKGKGNKAAQPGSGKGKVQFQGKKTKFASDDEHDEHDENGATGPVKRAREETDKEEPASKQQKTENGAGDQ |
| 5 | ANFLVHSSNNFGAILSSTNVGSNTYX | MPLNVSFTNRNYDLDYDSVQPYFYCDEEENFYQQQQQSELQPPAPSEDIWKKFELLPTPPLSPSRRSGLCSPSYVAVTPFSLRGDNDG |
| 6 | SNQNNFVHDCVNITIKQHTVTTTTKGENFTETDVKMMERVVEQMCITQYERESQAYYQRG | EPVRKEPVFTLAELVNDITPENLHENIDWGEPKDKEVW |
| 7 | GVVAAAEKTKQGVAEAAGKTKEGVLYVGSKTKEGVVHGVATVAEKTKEQVTNVGGAVVTGVTAVAQKTVEGAGSIAAATGFVKK | MGRRFLVTVRIQRAGRPLQERVFLVKFVRSRRPRTAS |
| 8 | QKLVFFAENVGSNKGAIIGLMVGGVV | VKSDGNILDDLNEATKKASDFVTDKTKEALADGEKTKDYIVEKTIEANETATEEAKKALDYVTEKGKEAGNKAAEFVEGKAEEAKNATKS |
| 9 | MNTLFNLFFDFITGILKNIGNIAA | HLLSFKKELGTLTSAINRRSTKQKKR |
| 10 | DQLETQTRELETAYSNLLRD | TTPPSVYPLAPGSAAQTNSMVTLGCLVKGYFPEPVTVTWNSGSLSSGVHTFPAVLQSDLYTLSSSVTVPSSTWPSETVTCNVAHPASSTKVDKKIVPR |

| Set 9 | Positive | Negative |
|---|---|---|
| 1 | SYGSSSQSSSYGQPQSGSYSQQPSYGGQQQSYGQQQSYN | MNTDAIESMVRDVLSRMNSLQGEAPAAAPAAGGASR |
| 2 | MNFGAFSINPAMMAAAQAALQSSWGMMGMLASQQNQ | MMFNKQIFTILILSLSLALAGSGCISEGAEDNVAQEI |
| 3 | GFGNSRGGGAGLGNNQGSNMGGGMNFGEFSIN | MKAAVDLKPTLTIIKTEKVDLELFPSPDMECADVPLLTPSSKEMMSQAL |
| 4 | ANFLVHSSNNFGAILSSTNVGSNTYX | YLDSGLGAPVPYPDPLEPKREVCELNPNCDELADHIGFQEAYQRFYGPV |
| 5 | GYGNQGGGYGGGYDNYGGGNYGSGNYNDFGNYNQQPSNYGPMKSGNFGGSRNMGGPY | LRKLNPPDESGPGCMSCKCVLS |
| 6 | RPLVNIYNCSGVQVGDNNYLTMQQTTALP | MSTESALSYAALILADSEIEISSEKLLTLTNAANVPDENIWADIFAKALDGQNLKDLLVNFSAGAAAPAGVAGGVAGGEAGEAEAEKEEEEAKEESDDDMGFGLFD |
| 7 | FLNCYVSGFHPSDIEVDLLKNGERIEKVEHSDLSFSKDWSFYLLYYTEFTPTEKDEYACRVNHV | PGDPDMIRYIDEFGQTTTRMQ |
| 8 | PLVNIYNCSGVQVGD | MSSENCFVAENSSLHPESGQENDATSPHFSTRHEGSFQVPVLCAVMNVVFITILIIALIALSVGQYNCPGQYTFSMPSDSHVSS |

| | | |
|---|---|---|
| 9 | KTKEGVVHGVATVAEKTKEQVTNVGGAVVTGVTAVAQKTVE | MMNNNGNQVSNLSNALRQVNIGSRNSNTTTDQSNINFEFSTGVNNNNNNNSSSNNNNVQNNNSGRNGSQNNDNENNIKNTLEQHRQQQQA |
| 10 | QKLVFFAEDVGSNKGAIIGLMVGGVVIA | RKRWQNEKLGLDAGDEYEDENLYEGLNLDDCSMYEDISRGLQGTYQDVGSLNIGDVQLEKP |

| Set 10 | Positive | Negative |
|---|---|---|
| 1 | GQRVTISCSGRSSNIGRNLVKWYQQFPGTAPKLLIYSNDQRPSGVPDRFSGSKSGTSASLAVSGLQSEDEADYYCAAWDATLNAWVFGGGT | MPVAGSELPRRPLPPAAQERDAEPRPP |
| 2 | SFFSFLGEAFDGARDMWRAYSDMREANYIGSDKYFHARGNYDAAKRGPGGVWAA | MDIKNSPSSLNSPSSYNCSQSILPLEHGSIYIPSSYVDSHHEYPAMTFYSPAVMNYSIPSNVTNLEGGPGRQTTSPNVLWPTPGHLSPLVVHRQLSHLYAEPQ |
| 3 | GYGNQGGGYGGGYDNYGGGNYGSGNYNDFGNYNQQPSNYGPMKSGNFGGSRNMGGPY | AKKAGAAKAKKPAGAAKKPKKATGTATPKKSTKKTPKKAKKPAAGAKKAKSPKKAKATKAKKAPKSPAKARAVKPKAAKPKTSKPKAAKPKKTAAKKK |
| 4 | RPLVNIYNCSGVQVGDNNYLTMQQTTALP | AGHELQPLAIVDQRPSSRASSRASSRPRPDDLEI |
| 5 | KVQIINKKLDLSNVQSKCGSKDNIKHVPGGGSVQIVYKPVDLSKVTSK | DGLVGMAIVGGMALGVAGLAGLIGLAVSKSKS |
| 6 | GGKVQIVYKPVDLSKVTSKCGSLGNIHH | MLSVRTPLATIADQQQLQLSPLKRLTLADKENTPPTLSSTRVLASKAARRIFQDSAELESKAPT |
| 7 | KTNMKHMAGAAAAGAVVGGLGGYMLGSAMSRPIIHFGSDY | MEEEKHHHHHLFHHKDKAEEGPVDYEKEIKHHKHLEQIGKLGTVAAGAYALHEKHEAKKDPEHAHKHKIEEEIAAAAAVGAGGFAFHEHHEKKDAKKEEKKKLRGDTTISSKLLF |
| 8 | PLVNIYNCSGVQVGD | VEPYSQEEAERAAGMGSYVPPRRLAVPTEVSTEVPEMDTST |
| 9 | KTKEGVVHGVATVAEKTKEQVTNVGGAVVTGVTAVAQKTVE | YNGQSETSQQPETPVFNTLPMMGKASPVSLGVPSEATANNGQQQQVQEQRRRINAMLQDYELQRRLHSEQLQFEQAQTQQAAVQVPGIQTLGTQSQ |
| 10 | MNTLFNLFFDFITGILKNIGNIAA | MLSLKKYLTEGLLQFTILLSLIGVRVDVDTYLTSQLPPLREIILGPSSAYTQTQFHNLRNTLDGYGIHPKSIDLDNYFTARRLLSQVRALDRFQVPTT |

Table S2: Definition of the different classification methods for amino acids groups

| Classification | group name | amino acids |
|---|---|---|
| 1 | A | R, K, E, D, Q, N |
| | B | G, A, S, T, P, H, Y |
| | C | C, V, L, I, M, F, W |
| | X | X |
| 2 | A | H, N, T, Q, C, S |
| | B | K, R, E, D |
| | C | I, L, M, V, W, Y, F, A |
| | G | G |
| | P | P |
| | X | X |
| 3 | A | H, T, C, S |
| | B | K, R, E, D |
| | C | I, L, M, V, W, Y, F, A |
| | D | Q, N |
| | G | G |
| | P | P |
| | X | X |

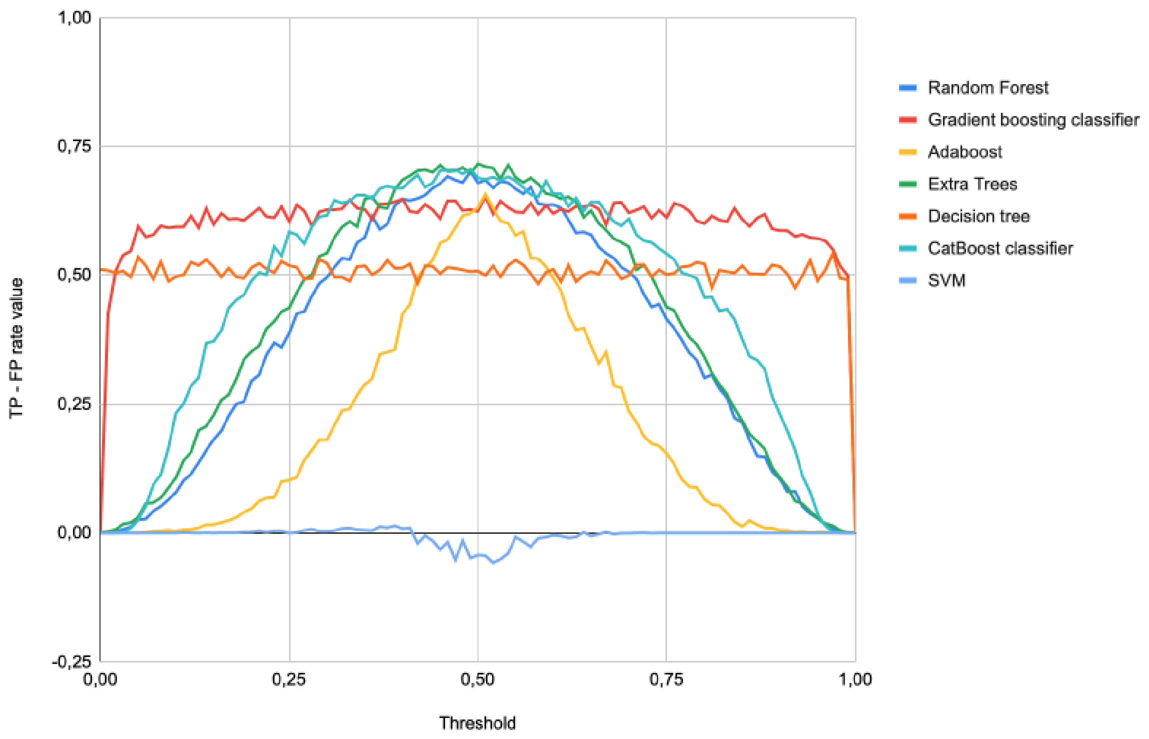

Figure S2: TP-FP rate comparison for a threshold going from 0 to 1 of the tested models.

#### Table S3: Paired comparison of the accuracy distribution of tested models

| Models | Random Forest | Gradient boosting | AdaBoost | Extra Trees | Decision tree | CatBoost classifier | SVM |
|---|---|---|---|---|---|---|---|
| Random Forest | | | | | | | |
| Gradient boosting | **** 2,252e-6 | | | | | | |
| AdaBoost | **** 6,974e-10 | N.S. 0,138 | | | | | |
| Extra Trees | N.S. 0,099 | **** 1,012e-9 | **** 7,252e-14 | | | | |
| Decision tree | **** 1,689e-48 | **** 1,837e-26 | **** 1,695e-20 | **** 1,426e-53 | | | |
| CatBoost classifier | N.S. 0,269 | **** 1,562e-8 | **** 1,742e-12 | N.S. 0,561 | **** 1,738e-51 | | |
| SVM | **** 1,986e-159 | **** 7,528e-153 | **** 3,298e-149 | **** 8,981e-161 | **** 1,937e-114 | **** 5,344e-158 | |

Wilcoxon-Mann-Whitney test result table. p-value > 0.05: N.S., p-value < 0.05: *, p-value < 0.01: **, p-value < 0.001: *** and p-value < 0.0001: ****.

Table S4: Results by test groups:

| Set 1 | ArchCandy2.0 | Tango | Pasta2.0 (Settings 1) | Pasta2.0 (Settings 2) | Pasta2.0 (Settings 3) | AmyloGram |
|---|---|---|---|---|---|---|
| Recall | 0.900 | 0.400 | 0.400 | 1 | 1 | 1 |
| Precision | 0.643 | 0.500 | 0.500 | 0.588 | 0.500 | 0.588 |
| Accuracy | 0.700 | 0.500 | 0.500 | 0.650 | 0.500 | 0.650 |
| F1 score | 0.750 | 0.444 | 0.444 | 0.741 | 0.667 | 0.741 |

| Set 2 | ArchCandy2.0 | Tango | Pasta2.0 (Settings 1) | Pasta2.0 (Settings 2) | Pasta2.0 (Settings 3) | AmyloGram |
|---|---|---|---|---|---|---|
| Recall | 0.900 | 0.300 | 0.200 | 0.500 | 0.800 | 0.600 |
| Precision | 0.692 | 0.600 | 0.667 | 0.500 | 0.500 | 0.545 |
| Accuracy | 0.750 | 0.550 | 0.550 | 0.500 | 0.500 | 0.550 |
| F1 score | 0.783 | 0.400 | 0.308 | 0.500 | 0.615 | 0.571 |

| Set 3 | ArchCandy2.0 | Tango | Pasta2.0 (Settings 1) | Pasta2.0 (Settings 2) | Pasta2.0 (Settings 3) | AmyloGram |
|---|---|---|---|---|---|---|
| Recall | 0.800 | 0.100 | 0.400 | 0.600 | 0.900 | 0.700 |
| Precision | 0.800 | 0.333 | 0.500 | 0.462 | 0.529 | 0.500 |
| Accuracy | 0.800 | 0.450 | 0.500 | 0.450 | 0.550 | 0.500 |
| F1 score | 0.800 | 0.154 | 0.444 | 0.522 | 0.667 | 0.583 |

| Set 4 | ArchCandy2.0 | Tango | Pasta2.0 (Settings 1) | Pasta2.0 (Settings 2) | Pasta2.0 (Settings 3) | AmyloGram |
|---|---|---|---|---|---|---|
| Recall | 1 | 0.300 | 0.300 | 0.500 | 0.700 | 0.600 |
| Precision | 0.714 | 0.600 | 0.429 | 0.455 | 0.500 | 0.462 |
| Accuracy | 0.800 | 0.550 | 0.450 | 0.450 | 0.500 | 0.450 |
| F1 score | 0.833 | 0.400 | 0.353 | 0.476 | 0.583 | 0.522 |

| Set 5 | ArchCandy2.0 | Tango | Pasta2.0 (Settings 1) | Pasta2.0 (Settings 2) | Pasta2.0 (Settings 3) | AmyloGram |
|---|---|---|---|---|---|---|
| Recall | 1 | 0.300 | 0.200 | 0.500 | 0.900 | 0.700 |
| Precision | 0.750 | 0.600 | 0.667 | 0.556 | 0.563 | 0.583 |
| Accuracy | 0.667 | 0.550 | 0.550 | 0.550 | 0.600 | 0.600 |
| F1 score | 0.800 | 0.400 | 0.308 | 0.526 | 0.692 | 0.636 |

| Set 6 | ArchCandy2.0 | Tango | Pasta2.0 (Settings 1) | Pasta2.0 (Settings 2) | Pasta2.0 (Settings 3) | AmyloGram |
|---|---|---|---|---|---|---|
| Recall | 0.800 | 0.200 | 0.200 | 0.600 | 0.900 | 0.700 |
| Precision | 0.667 | 0.667 | 1 | 0.750 | 0.563 | 0.700 |
| Accuracy | 0.700 | 0.550 | 0.600 | 0.857 | 0.600 | 0.700 |
| F1 score | 0.727 | 0.308 | 0.333 | 0.706 | 0.692 | 0.700 |

| Set 7 | ArchCandy2.0 | Tango | Pasta2.0 (Settings 1) | Pasta2.0 (Settings 2) | Pasta2.0 (Settings 3) | AmyloGram |
|---|---|---|---|---|---|---|
| Recall | 0.800 | 0.400 | 0.500 | 0.800 | 1 | 0.900 |
| Precision | 0.615 | 0.667 | 0.625 | 0.500 | 0.500 | 0.529 |
| Accuracy | 0.650 | 0.600 | 0.600 | 0.500 | 0.500 | 0.550 |
| F1 score | 0.696 | 0.500 | 0.556 | 0.615 | 0.667 | 0.667 |

| Set 8 | ArchCandy2.0 | Tango | Pasta2.0 (Settings 1) | Pasta2.0 (Settings 2) | Pasta2.0 (Settings 3) | AmyloGram |
|---|---|---|---|---|---|---|
| Recall | 0.900 | 0.400 | 0.400 | 0.600 | 0.900 | 0.700 |
| Precision | 0.750 | 0.667 | 0.667 | 0.462 | 0.500 | 0.583 |
| Accuracy | 0.800 | 0.600 | 0.600 | 0.450 | 0.500 | 0.600 |
| F1 score | 0.818 | 0.500 | 0.667 | 0.522 | 0.643 | 0.636 |

| Set 9 | ArchCandy2.0 | Tango | Pasta2.0 (Settings 1) | Pasta2.0 (Settings 2) | Pasta2.0 (Settings 3) | AmyloGram |
|---|---|---|---|---|---|---|
| Recall | 0.800 | 0.300 | 0.400 | 0.600 | 0.800 | 0.900 |
| Precision | 0.571 | 0.600 | 0.667 | 0.429 | 0.444 | 0.600 |
| Accuracy | 0.600 | 0.550 | 0.600 | 0.400 | 0.400 | 0.650 |
| F1 score | 0.667 | 0.400 | 0.667 | 0.500 | 0.571 | 0.636 |

| Set 10 | ArchCandy2.0 | Tango | Pasta2.0 (Settings 1) | Pasta2.0 (Settings 2) | Pasta2.0 (Settings 3) | AmyloGram |
|---|---|---|---|---|---|---|
| Recall | 0.800 | 0.400 | 0.400 | 0.800 | 0.900 | 0.900 |
| Precision | 0.615 | 0.667 | 0.667 | 0.571 | 0.529 | 0.600 |
| Accuracy | 0.650 | 0.600 | 0.600 | 0.600 | 0.550 | 0.650 |
| F1 score | 0.696 | 0.500 | 0.667 | 0.667 | 0.667 | 0.720 |

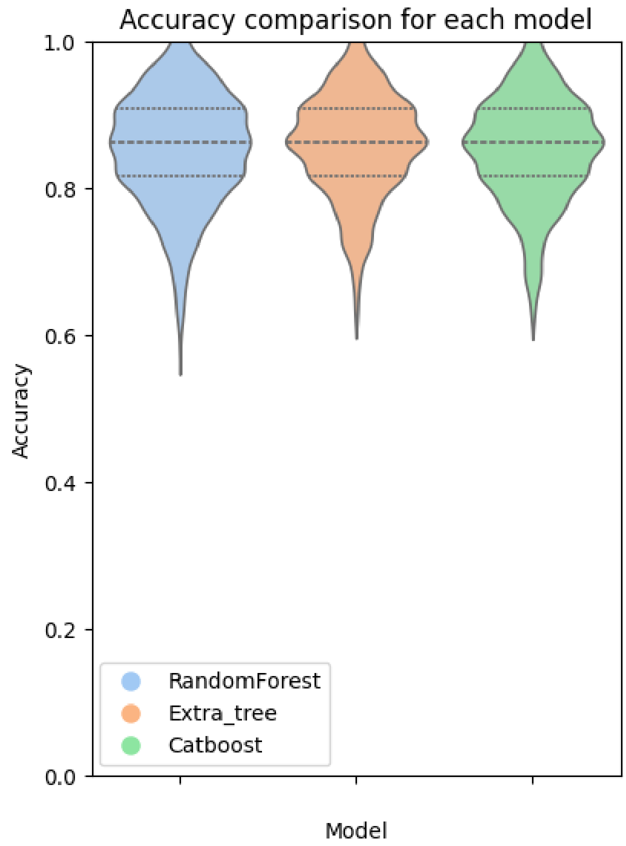

Figure S3: Accuracy comparison of optimized Random Forest, Extra trees and CatBoost classifiers
